# Epigenetic divergence is sufficient to trigger heterosis in *Arabidopsis thaliana*

**DOI:** 10.1101/059980

**Authors:** Kathrin Lauss, René Wardenaar, Marieke H.A. van Hulten, Victor Guryev, Joost J.B Keurentjes, Maike Stam, Frank Johannes

## Abstract

Despite the importance and wide exploitation of heterosis in commercial crop breeding, the molecular mechanisms behind this phenomenon are not well understood. Interestingly, there is growing evidence that beside genetic also epigenetic factors contribute to heterosis. Here we used near-isogenic but epigenetically divergent parents to create epigenetic F1 hybrids (epiHybrids) in Arabidopsis, allowing us to quantify the contribution of epigenetics to heterosis. We measured traits such as leaf area (LA), growth rate (GR), flowering time (FT), main stem branching (MSB), rosette branching (RB) and final plant height (HT) and observed several strong positive and negative heterotic phenotypes among the epiHybrids. For LA and HT mainly positive heterosis was observed, while FT and MSB mostly displayed negative heterosis. Heterosis for FT, LA and HT could be associated with several heritable, differentially methylated regions (DMRs) in the parental genomes. These DMRs contain 35 (FT and LA) and 14 (HT) genes, which may underlie the heterotic phenotypes observed. In conclusion, our study indicates that epigenetic divergence can be sufficient to cause heterosis.

**Author Summary:** Crossing two genetically distinct parents generates hybrid offspring. Sometimes hybrids are performing better than their parents in particular traits and this is referred to as heterosis. Hybridization and heterosis are naturally occurring processes and crop breeders intentionally cross genetically different parental lines in order to generate hybrids with maximized traits such as yield or stress tolerance. So far, the mechanisms behind heterosis are not well understood. In this study we focused on the effect of epigenetic variation onto heterosis in hybrids, and for this purpose we created epigenetic hybrids (epiHybrids) by crossing wildtype plants with a selection of genetically very similar but epigenetically divergent lines. An extensive phenotypic analysis of the epiHybrids and their parental lines showed that epigenetic divergence between parental genomes can be a major determinant of heterosis. Importantly, multiple heterotic phenotypes could be associated with meiotically heritable differentially methylated regions (DMRs) in the parental genomes, allowing us to map epigenetic quantitative trait loci (QTLs) for heterosis. Our results indicate that epigenetic variation can contribute to heterosis and suggests that heritable epigenetic variation could be exploited for the improvement of crop traits.

## Introduction

Heterosis describes an F1 hybrid phenotype that is superior compared to the phenotype of its parent varieties. The phenomenon has been exploited extensively in agricultural breeding for decades and has improved crop performance tremendously [1,2]. Despite its commercial impact, knowledge of the molecular basis underlying heterosis remains incomplete. Most studies mainly focused on finding genetic explanations, resulting in the classical dominance [1,3,4] and overdominance [4,5] models describing heterosis. In line with genetic explanations it has been observed that interspecies hybrids often show a higher degree of heterosis than intraspecies hybrids, indicating that genetic distance correlates with the extent of heterosis [2,6]. However, genetic explanations do often not sufficiently explain nor predict heterosis. There is growing evidence that also epigenetic divergence plays a role in heterosis [7–9]. It has, for example, been shown that altered epigenetic profiles at genes regulating circadian rhythm play an important role in heterotic Arabidopsis hybrids [10]. Moreover, heterotic hybrids of Arabidopsis, maize and tomato are shown to differ in levels of small regulatory RNAs and/or DNA methylation (5mC) relative to their parental lines [11–14]. Processes such as the transfer of 5mC between alleles (trans chromosomal methylation, TCM), or a loss of 5mC at one of the alleles (trans chromosomal demethylation, TCdM) have been indicated to contribute to the observed remodeling of the epigenome [7,13,15]. Strikingly, some of these changes in 5mC levels have been shown to be stable over multiple generations [15,16].

In this study, we demonstrate that heterotic phenotypes occur in *A. thaliana* F1 epigenetic hybrids (epiHybrids) that were generated from near-isogenic but epigenetically very divergent parental lines. Moreover, we found that some of those heterotic phenotypes could be associated with differentially methylated regions (DMRs) in their parental genomes, allowing us to map QTLs for heterosis.

## Results and Discussion

### Construction of epigenetic Hybrids

Hybrids are usually generated from parental lines that vary at both the genomic and epigenomic level and disentangling those two sources of variation is challenging. To overcome this limitation, we generated epigenetic *A. thaliana* F1 hybrids (epiHybrids) from near-isogenic but epigenetically divergent parental lines by crossing Col-0 wildtype (Col-wt) as maternal parent to 19 near-isogenic *ddm1-2*-derived epigenetic recombinant inbred lines (epiRILs) [17] as the paternal parents (Fig 1a).

**Fig 1.**
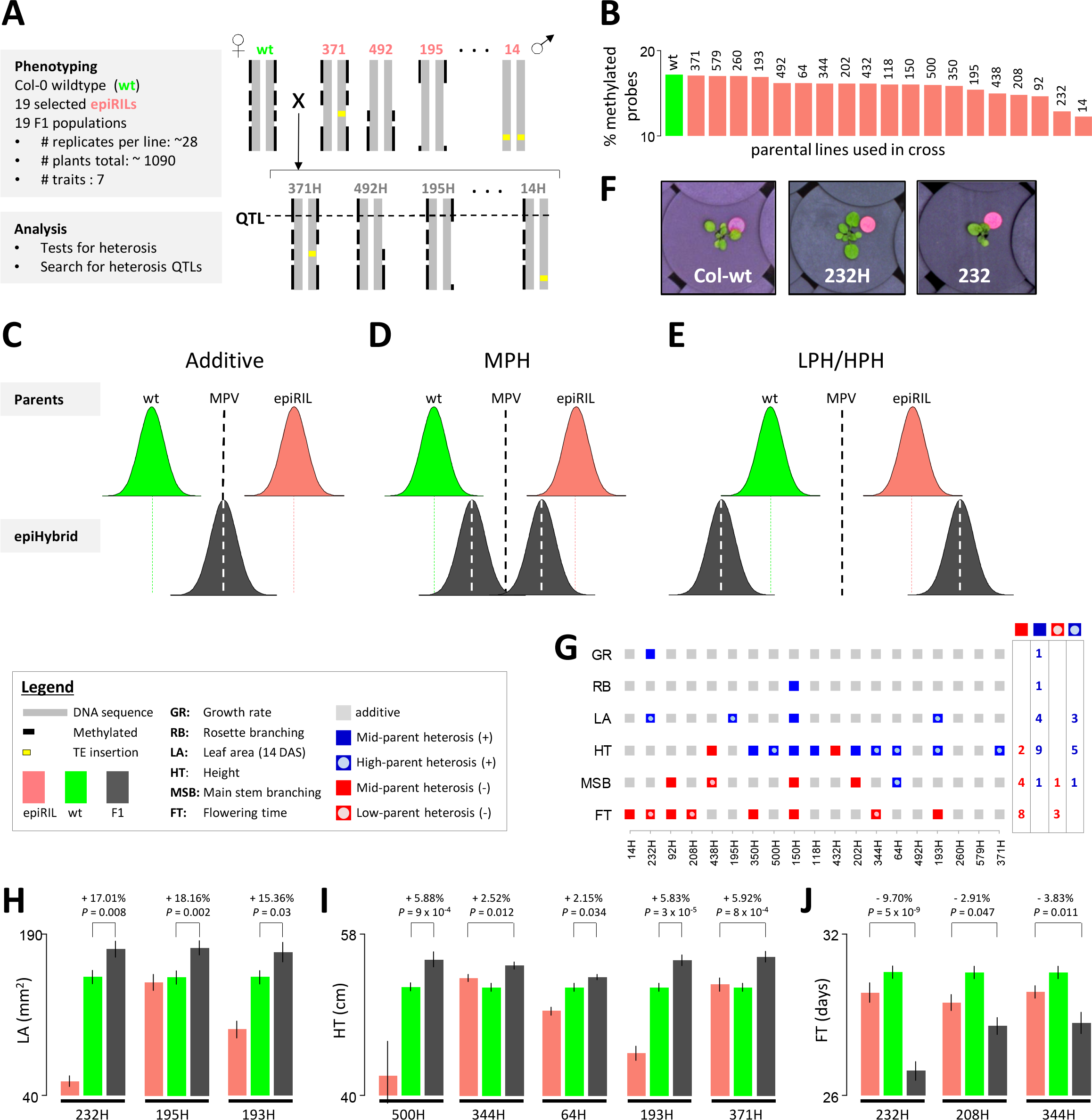
Heterosis occurs in epiHybrids. (**a**) Experimental setup. Lines are depicted schematically as one chromosome with the numbers indicating the epiRIL ID (e.g. 371 & 492) and the respective epiHybrid (e.g. 371H & 492H). (**b**) Genome-wide 5mC levels (y-axis) of the Col-wt line in green and the epiRIL parental lines in salmon. Numbers indicate the epiRIL IDs. The 5mC levels were calculated as the proportion of methylated MeDIP probes with respect to the total amount of probes. **(c-e)** Three classes of phenotypic effects monitored in the epiHybrids. The black dashed line indicates the mid-parent value. The green and salmon dashed lines indicate the mean performance of the parental lines. The white dashed lines indicate the mean performance of the epiHybrids. **(f)** Col-wt, epiHybrid 232H and epiRIL 232 at 13 days after sowing as an example for high-parent heterosis. **(g)** Phenotypic effects in six traits monitored across the 19 epiHybrids. The right panel summarizes positive and negative heterotic effects per trait. **(h-j)** Examples of epiHybrids exhibiting high-parent heterosis in leaf area and height (LA and HT; **h** and **i**), and low-parent heterosis in flowering time (FT; **j**) Error bars, ± 1 SEM. Deviation from high parent or low parent is shown in percent.

DDM1 (*DECREASE IN DNA METHYLATION 1*) is a nucleosome remodeler and a *ddm1-2* deficiency leads to a severe loss of 5mC [18], primarily in long transposable elements and other repeat sequences [19]. EpiRILs carry chromosomes that are a mosaic of Col-wt and hypomethylated *ddm1-2*-derived genomic regions [17,20,21] (Fig 1a). Nineteen epiRIL parental lines were selected that sample a broad range of 5mC divergence from the Col-wt reference methylome (Fig 1b, S1 Table). Besides, lines were chosen that have a wildtype methylation profile at *FWA* (S1 Fig, S1 Table), as loss of DNA methylation at the *FWA(FLOWERING LOCUS WAGENINGEN)* locus is known to affect flowering time [22]. Furthermore, we selected for a range of phenotypic variation in two traits that have previously been monitored in the epiRILs, flowering time and root length (S1 Table); outliers were excluded [17]. With our experimental design we could demonstrate, as proof-of-principle, the extent to which divergence in 5mC profiles in parental lines can contribute to heterosis.

### Heterotic phenotypes occur in the epiHybrids

The phenotypic performance of the 19 epiHybrids and their parental lines was assessed by monitoring about 1090 plants (~28 replicates per line) for a range of quantitative traits: LA, GR, FT, MSB, RB, HT and SY (S2-S7 Tables). The phenotypic observations for SY were inconsistent in a replication experiment, therefore those datasets were excluded from further analysis. The hybrids and parental lines were grown in parallel in a climate-controlled chamber with automated watering. The plants were randomized throughout the chamber to level out phenotypic effects caused by plant position. LA was measured up to 14 days after sowing (DAS), using an automated camera system (Fig 1f), and growth rate (GR) was determined based on this data (SI text). FT was scored manually as opening of the first flower. After all plants started flowering, the plants were transferred to the greenhouse and grown to maturity. MSB, RB and HT were scored manually after harvesting of the plants.

The extent of heterosis was evaluated by comparing the hybrid performance with its parental lines. We distinguished five effects (Fig 1c-e): additivity, positive mid-parent heterosis (positive MPH), negative mid-parent heterosis (negative MPH), high-parent heterosis (HPH) and low-parent heterosis (LPH). An additive effect describes a hybrid performance that is equal or close to the average performance of the two parents (the mid-parent value, MPV). MPH refers to deviations in percent from the MPV in positive or negative direction. Hybrids displaying MPH are further tested for HPH and LPH, which describe hybrid performance exceeding the high parent, or falling below the lowest parent, respectively. In crop breeding, the focus is usually on obtaining HPH and LPH as these present novel phenotypes that are outside the parental range. Depending on the trait monitored and commercial application, either HPH or LPH can be considered superior. For instance, early flowering may be preferable over late flowering; in such cases maximizing LPH may be desirable. For other traits, such as yield or biomass, it is more important to maximize HPH. However, in order to obtain a comprehensive view of hybrid performance it is informative to also track MPH in addition to LPH and HPH, because many mature traits may be affected by other traits that do not display fully penetrant heterotic effects.

We observed a remarkably wide range of heterotic phenotypes among the epiHybrids (Fig 1g, S2-19 Tables). The magnitude of these phenotypic effects was substantial (Fig 1h-j, S2 Fig, S8-19 Tables) and similar to that typically seen in hybrids of Arabidopsis natural accessions[23,24]. Many epiHybrids (16/19) exhibited significant MPH in at least one of the six monitored traits (FDR = 0.05, Fig 1g). Across all hybrids and traits, we observed 30 cases of positive MPH and negative MPH. Among those, four cases show LPH and nine cases show HPH (Fig 1g). Interestingly, in 11 out of the 17 cases of MPH the phenotypic means of the epiHybrids were in the direction of the phenotypic means of the epiRIL parent rather than in the direction of the Col-wt parent (S2-7 Tables, F1 trend). Also all four LPH and two of the HPH cases were in the direction of the epiRIL parent (Fig 1i-j, S2 Fig). This observation illustrates that *ddm1-2*-derived hypomethylated epialleles are often (partially) dominant over wild-type epialleles, which contrasts the situation seen in EMS screens where novel mutations typically act recessively.

We observed cases of HPH for LA, HT and MSB, and cases of LPH for FT and MSB. HPH for LA occurred in epiHybrids 232H, 195H and 193H (3/19 epiHybrids). Those epiHybrids significantly exceeded their best parent (Col-wt) by 17%, 18% and 15%, respectively (Fig 1h, S19 Table). Interestingly, although growth rate (GR) is developmentally related to LA, hybrid effects in GR were only moderately, albeit positively, correlated with LA (rho = 0.57, P = 0. 02), which implies that LA heterosis is determined by other traits besides GR.

For HT we detected five cases of significant HPH with up to 6% increases in HT (Fig 1i, S14 Table). One may expect LA HPH to strongly correlate with HT HPH, as the rosette is providing nutrients for the developing shoot[25]. However, HPH for both LA and HT occurred only in one epiHybrid (193H; Fig 1g).

For MSB, we detected one case of HPH (64H; Fig 1g and S2 Fig).

Besides positive heterosis, our phenotypic screen revealed strong negative heterotic effects for FT (earlier flowering) and MSB (less main stem branching). Significant LPH occurred in the epiHybrids 232H, 208H and 344H (FT) and 438H (MSB) (Fig 1j, S2 Fig, S15 and S17 Tables). In the most prominent case for FT (232H), FT was about 10% earlier than that of the earliest flowering parent. 208H and 244H flowered 3% and 4% earlier than their lowest parent (epiRIL 208 and epiRIL 344), respectively. 438H showed 14% less MSB than the lowest parent (S2 Fig).

The reproducibility of our findings was tested by performing replicate experiments, using seeds from newly performed crosses and the same climate controlled growth chamber as before. We focused on epiHybrids that exhibited relatively strong positive or negative heterotic phenotypes in the initial screen (193H, 150H, 232H; Fig 1g), and measured LA, FT and HT. We found that the direction of the heterotic effects in LA, FT and HT was reproducible in all cases tested (Fig 2a and b). Importantly, the LA and HT HPH observed for 193H, and the strong FT LPH for 232H were perfectly reproducible, while LA HPH observed for 232H became positive MPH (Fig 2a). Taken together, these results show that the heterotic effects observed in the epiHybrids are relatively stable for LA, HT and FT, even across fresh parental seed batches and independently performed crosses, which is not always the case for Arabidopsis phenotypes [26].

**Fig 2.**
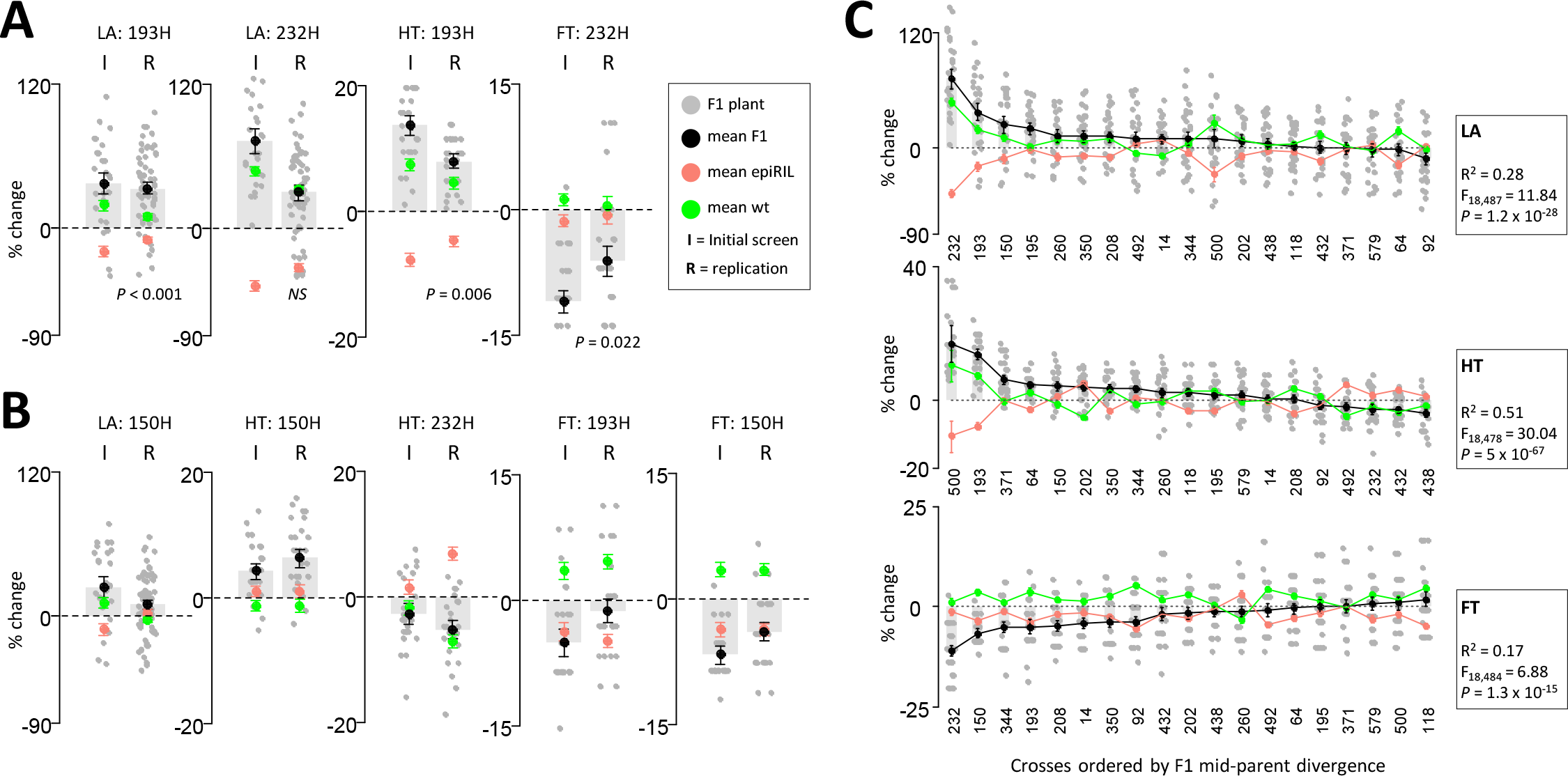
Confirmation of mid-parent (MP) divergence in the initial screen and replicate experiment for epiHybrids 150H, 193H and 232H. **(a)** Results for cases of HPH and LPH for LA, HT and FT in initial experiment. **(b)** Results for traits showing less eminent phenotypic effects for LA, HT and FT. The mid-parent value (MPV) is shown as a dashed horizontal line and the MP divergence is shown as change from MPV in percent. To illustrate the F1 epiHybrid distribution for each trait, the individual replicate plants are depicted as dots. **(c)** F1 MP divergence for LA, HT and FT for all epiHybrids. The MPV is shown as a horizontal dashed line and MP divergence is shown as change from MPV in percent. The epiHybrids are ordered from highest (left) to lowest (right) F1 MP divergence. To illustrate the F1 epiHybrid distribution for each trait, the individual replicate plants are depicted as dots. Variance component analysis was used to estimate how much of the total variation in MP divergence can be explained by between-cross variation. The F-statistic from this analysis is shown in the boxes.

### Heterotic phenotypes are associated with QTLs

To understand the sources of the LA, HT and FT heterotic effects observed among the ~530 epiHybrid plants, we calculated the phenotypic divergence of each epiHybrid plant from its respective mid-parent value. Using variance component analysis we estimated that 17%, 28% and 51% of the total variation in mid-parent divergence for FT, LA and HT, respectively, can be attributed to (epi)genomic differences between the Col-wt and epiRILs used for the crosses (Fig 2c, S20 Table, SI text). Global 5mC divergence between the Col-wt and the epiRILs parental lines could not account for this variation (S3 Fig). We therefore reasoned that heterotic phenotypes are due to (partial) dominance effects caused by specific regions being epi-heterozygous for an epiRIL-inherited hypomethylated epiallele (*U*) and a Col-wt-inherited methylated epiallele (*M*). To test this possibility, we used the methylomes of Col-wt and the epiRIL parents[20] to predict epi-homozygous (*MM*) and epi-heterozygous (*MU*) regions in the genomes of the epiHybrids (Fig 3a, SI text), and assessed whether heritable epigenetic differences at specific loci could explain the variation in MPH among crosses (S4 Fig). The analysis revealed two QTLs on chromosome (chr) 3 contributing to the between-cross variation in MPH in FT (QTL 1: LOD=3.12, 37.62 cM; QTL 2: LOD=3.33, 101.44 cM, Fig 3b; S21 Table). EpiHybrids epi-heterozygous (*MU*) at these loci showed significant negative MPH compared to their epi-homozygous (*MM*) counterparts (Fig 3c). While not significant at the genome-wide scale (Fig 3b), the same two QTLs had substantial suggestive effects on LA heterosis in the opposite direction than FT (Fig 3b and c), indicating that both QTLs act pleiotropically.

**Fig 3.**
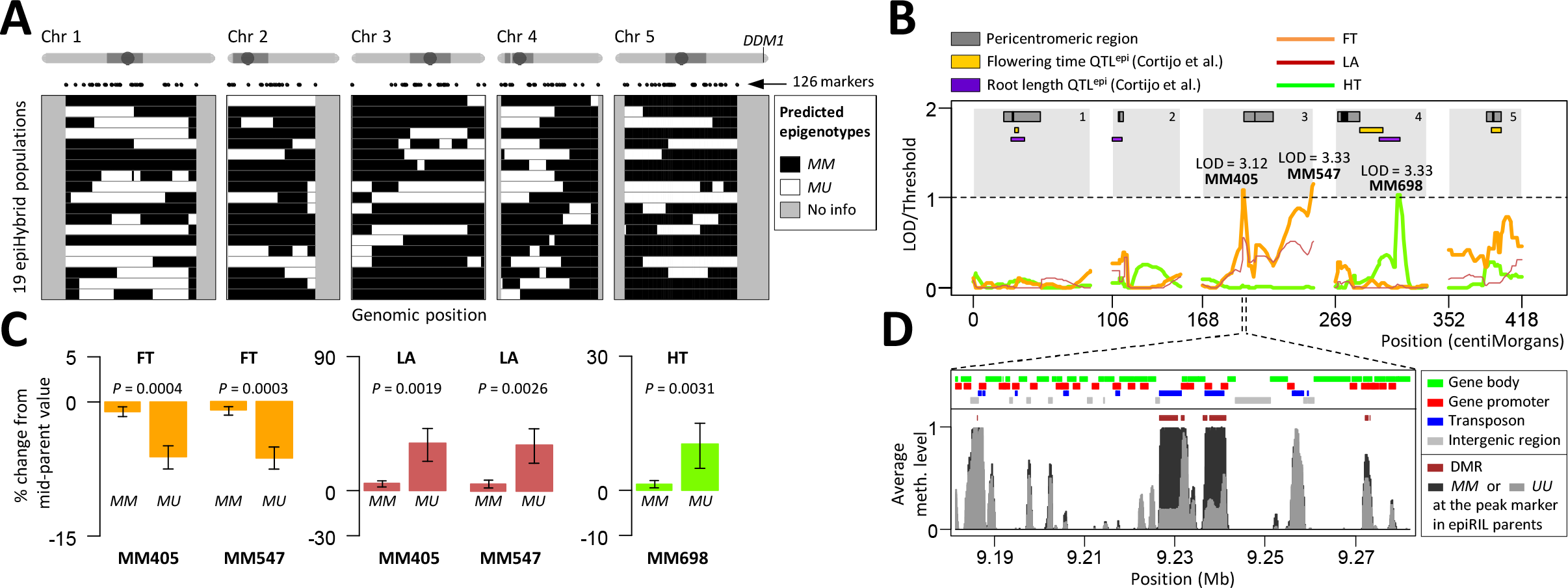
Interval mapping approach detects significant QTLs for mid-parent divergence. **(a)** Genome-wide patterns of Col-wt and *ddm1-2* inherited epi-haplotypes in the (epi)genomes of the parental epiRILs used in this study. **(b)** QTL profiles for FT, HT and LA. Published QTLs^epi^ for root length and flowering time are shown. **(c)** Effect direction of the QTLs. Error bars, ± 1 SE of the Estimate (SEE). **(d)** Zoom in of one of the QTL intervals of FT. The top panel shows the annotations along the genome. The bottom panel shows the locations of candidate DMRs and the average methylation level along the genome for epiRIL parents that are either methylated (*MM*) or unmethylated (*UU*) at the peak marker.

We also detected a single QTL locus on chr 4 (LOD=3.33, 56.00 cM) that contributes to the between-cross variation in MPH for HT (Fig 3b, S21 Table). In this case, *MU* epiHybrids showed significant positive MPH compared to *MM* epiHybrids (Fig 3c). Interestingly, the HT QTL overlaps with a previously identified QTL^epi^ for root length in the epiRILs[21]. The same study identified QTLs^epi^ associated with FT [21] that we did not detect here (Fig 3b), implying that different regions may play a role in FT trait variation than in FT heterosis.

### Heterotic phenotypes are associated with DMRs in the parental genomes

The detection of heterosis QTLs for FT, LA and HT provided a rationale to search for causal variants in the QTL confidence intervals. TE-associated structural variants (TEASVs) are known to occur at low frequency in a *ddm1-2*-derived DNA hypomethylated background [17,21,27,28], hence we re-analyzed whole-genome sequencing data from the epiRIL parents [21] for TEASVs but did not detect any that could account for the QTL effects, suggesting that the QTLs most likely have an epigenetic basis (SI text). Indeed, a thorough analysis of the methylomes of the parental epiRILs, using the available MeDIP tiling array data [20], identified 55 and 18 potentially causal differentially methylated regions (DMRs) in the FT, LA and HT QTL regions, mapping to 35 and 14 unique genes, respectively (Fig 3d, S5-S9 Figs, S22-S26 Tables, SI text). Potentially interesting genes in the candidate regions of the FT/LA QTLs (S25 Table) include for example RPL5A, which was shown to affect development through regulating auxin and influencing leaf shape and patterning [29,30], and AT3G26480, a protein that shows partial homology to GTS1, which has been implemented in biomass accumulation [31]. Another potentially interesting candidate is Chup1, which is crucial for chloroplast movement in leaves in response to light [32]. These candidate genes provide excellent targets for follow-up studies.

### Conclusions

In a recently published study, heterosis for rosette area was reported in an epigenetic F1 hybrid generated by crossing a *met1*-derived epiRIL with Col-wt [33]. *DNA-METHYLTRANSFERASE1 (MET1)* is involved in maintenance of DNA methylation at cytosines in CG sequence context and a mutation in this gene causes a severe loss of DNA methylation in the CG and CHH context [34]. Heterosis was observed in a parent-of origin manner; the reciprocal cross did not result in heterosis [33]. This suggests that the heterosis detected may be due to an effect of the maternal cytoplasm rather than differences in epigenetic marks in the parental genomes. Here, we used Col-wt as maternal parent in all crosses to specifically monitor phenotypic effects associated with the epiRIL methylomes. We observed a wide range of heterotic effects, and our proof-of-principle QTL mapping approach indicated that these phenotypic effects are very likely attributable to methylation differences between Col-wt and the epiRILs. Moreover, our results, together with those of Dapp et al. [9], indicate that heterosis in F1 hybrids generated from epigenetically divergent lines may be a more general phenomenon. A more recent study described widespread DNA methylation changes in an epiHybrid derived from Col-wt and a *met1*-mutant [34]. Remarkably, the formation of spontaneous non-parental epialleles was observed in the epiHybrid, mostly at pericentromeric transposon sequences, but also at genic loci [34]. This demonstrates that novel epigenetic variation, which is not readily predictable from the parental methylomes, can be created during hybridization. Future research needs to address if and how these methylome changes relate to phenotypic variation. This study also stresses that for a refined understanding of the effect of epigenetic QTLs as described in this study, methylation changes should be thoroughly analyzed.

## Material and Methods

### Plant Material

The epigenetic recombinant inbred lines (epiRILS) in our study were generated by Johannes et al [17].The epiRILs were constructed as follows: An *Arabidopsis thaliana* Col-0 line deficient for *ddm1-2 (DECREASE IN DNA METHYLATION 1)* was crossed to an isogenic Col-0 wildtype line (Col-wt) and the resulting F1 was backcrossed as female parent to Col-wt. Subsequently about 500 progeny plants with a wildtype *DDM1* allele were selected and propagated through six more rounds of selfing, generating a population of 500 different epiRILs. We selected 19 different epiRILs as paternal plants for generating epiHybrids (Line IDs: 14, 232, 92, 208, 438, 195, 350, 500, 150, 118, 432, 202, 344, 64, 193, 508, 260, 579, 371). Our selection criteria were as follows: 1) Wide range of DNA methylation divergence from Col-wt and among the selected lines; 2) Wildtype DNA methylation state at the FWA locus in order to avoid that differences in DNA methylation at this locus give rise to differences in flowering time [22] in the hybrids; 3) Wide range of phenotypic variation in flowering time and root length among the selected lines. The epiRIL lines were purchased from the Arabidopsis Stock center of INRA Versailles (http://publiclines.versailles.inra.fr/).

### Crosses

To generate F1 hybrids from the selected epiRIL lines and Col-wt, all parental plants were grown in parallel in soil (Jongkind 7 from Jongkind BV, http://www.jongkind.com/) in pots (Danish size 40 cell, Desch Plantpak, http://www.desch-plantpak.com/en/Home.aspx). The plants were grown at 20°C, 60% humidity, in long day conditions (16h light, 8h dark), and were watered 3 times per week. All crosses were performed in parallel in a time frame of two weeks to avoid phenotypic effects in the F1 progeny due to differences in growing conditions. To exclude that differences in maternal cytoplasm affect the phenotypes of the F1 plants, Col-wt plants were used as a maternal parent and the epiRILs as paternal parents. In parallel, all parental lines, Col-wt and epiRILS, were propagated by manual selfing. This to 1) ensure that parental and F1 hybrid seeds were generated under the same growing conditions and 2) exclude potential phenotypic effects derived from hand pollination[35].

### Phenotypic Screen

The seeds were stratified at 4°C for 3 days on petri-dishes containing filter paper and water before transferring them onto Rockwool/Grodan blocks (soaked in Hyponex NPK: 6.5 – 6.19 medium) in a climate controlled chamber (20°C, 70% humidity, long day conditions (16h light, 8h dark)). The transfer of the seeds onto the Rockwool blocks is defined as time point 0 days after sowing (DAS). Seeds from each parental and hybrid line were sown in 28 replicates and their positions were randomized throughout the growth chamber to level out phenotypic effects caused by plant position. The plants were watered two or three times per week depending on their size. After the plants started flowering, they were transferred to the greenhouse (20°C, 60% humidity, long day conditions (16h light, 8h dark)). In the greenhouse, the plants were watered 3 times per week and stabilized by binding them to wooden sticks at later developmental stages. The plants were harvested once the siliques of the main inflorescence and its side branches were ripe.

Rosette Leaf Area (LA): LA was monitored by an automated camera system (Open Pheno System, WUR) from 4 days after sowing (DAS). The system consists of 14 fixed cameras that can take pictures of up to 2145 plants daily, every two hours. We monitored LA until 14 DAS since at later time points leaves start overlapping hampering the correct detection of LA. Leaf area in mm2 was calculated by an ImageJ based measurement setup (http://edepot.wur.nl/169770).

Flowering time (FT): FT was defined as the DAS at which the first flower opened. FT was scored manually each day before 12am.

Height (HT): HT was scored manually in cm on dried plants. The measurement was taken at the main inflorescence, from the rosette to the highest flowerhead.

Branching: Branching was scored on the dried plants by counting the branches emerging from the rosette (RB) and from the main stem (MSB).

Total Seed Yield (SY): Seeds were harvested from the dried plants, cleaned by filtering and seed yield was subsequently determined by weighing (resulting in mg seeds per plant).

#### Data analysis

For the data analysis see the Supplementary Information.

### Replication experiment with selected hybrids

Freshly ordered seeds of epiRILs (Line IDs: 92, 150, 193, 232) from the Arabidopsis Stock center Versailles were used for the replication experiment with the hybrids selected. The crosses with the epiRILS and the phenotypic screen were performed as described above with the exception that more replicates were monitored for each parental and hybrid line: 60 replicates for LA and 30 replicates for the traits FT and HT. Furthermore, branching was not examined in the replication experiment.

## Acknowledgements

This work was supported by the Centre for Improving Plant Yield (CIPY)(part of the Netherlands Genomics Initiative and the Netherlands Organization for Scientific Research). We thank F. Becker, I. HÖvel, D. Angorro, R. Kooke, J.A. Bac-Molenaar, M. Tark-Dame, P. Sanderson, M. Koini, T. Bey, B. Weber, L. Tikovsky and Unifarm Wageningen for technical support during sowing or phenotyping. We thank H. Westerhoff for discussion and critically reading the manuscript.

## Author Contributions

K.L., M.S. and F.J. designed the study, interpreted the data and wrote the manuscript with contributions from J.J.B.K. and R.W. K.L. and M.H.A.v.H. planned and performed the phenotypic screen. F.J. and R.W. performed the data analysis. V.G. analyzed sequencing data of the epiRIL parents.

## References

1. Schnable PS, Springer NM. Progress toward understanding heterosis in crop plants. Annu Rev Plant Biol. 2013;64: 71–88. doi:10.1146/annurev-arplant-042110-103827

2. Chen ZJ. Molecular mechanisms of polyploidy and hybrid vigor. Trends Plant Sci. 2010;15: 57–71. doi:10.1016/j.tplants.2009.12.003

3. Jones DF. Dominance of linked Factors as a means of accounting for Heterosis. Genetics. 1917;2: 466–479.

4. Crow JF. Anectdotal, Historical and Critical Commentaries on Genetics 90 Years Ago: The Beginning of Hybrid Maize. Genetics. 1998;148: 923–928.

5. Crow JF. Alternative Hypotheses of Hybrid Vigor. Genetics. 1948;33: 477–487. Available: http://www.pubmedcentral.nih.gov/articlerender.fcgi?artid=1209419&tool=pmcentrez&rendertype=abstract

6. East EM. Heterosis. Genetics. 1936;21: 336–397.

7. Groszmann M, Greaves IK, Fujimoto R, James Peacock W, Dennis ES. The role of epigenetics in hybrid vigour. Trends Genet. 2013;29: 684–690. Available: http://linkinghub.elsevier.com/retrieve/pii/S016895251300125X

8. Springer NM. Epigenetics and crop improvement. Trends Genet. Elsevier Ltd; 2013;29: 241–247. doi:10.1016/j.tig.2012.10.009

9. Dapp M, Reinders J, Bediee A, Balsera C, Bucher E, Theiler G, et al. Heterosis and inbreeding depression of epigenetic Arabidopsis hybrids. Nat Plants. 2015;1: 1–8. doi:10.1038/nplants.2015.92

10. Ni Z, Kim E-D, Ha M, Lackey E, Liu J, Zhang Y, et al. Altered circadian rhythms regulate growth vigour in hybrids and allopolyploids. Nature. 2009;457: 327–333. doi:10.1038/nature07523

11. Groszmann M, Greaves IK, Albertyn ZI, Scofield GN, Peacock WJ, Dennis ES. Changes in 24-nt siRNA levels in Arabidopsis hybrids suggest an epigenetic contribution to hybrid vigor. Proc Natl Acad Sci U S A. 2011;108: 2617–2622. Available: http://eutils.ncbi.nlm.nih.gov/entrez/eutils/elink.fcgi?dbfrom=pubmed&id=21266545&retmode=ref&cmd=prlinks

12. Barber WT, Zhang W, Win H, Varala KK, Dorweiler JE, Hudson ME, et al. Repeat associated small RNAs vary among parents and following hybridization in maize. Proc Natl Acad Sci U S A; 2012;109: 10444–10449. Available: http://eutils.ncbi.nlm.nih.gov/entrez/eutils/elink.fcgi?dbfrom=pubmed&id=22689990&retmode=ref&cmd=prlinks

13. Shivaprasad P V, Dunn RM, Santos BA, Bassett A, Baulcombe DC. Extraordinary transgressive phenotypes of hybrid tomato are influenced by epigenetics and small silencing RNAs. EMBO J.; 2012;31: 257–266. doi:10.1038/emboj.2011.458

14. Shen H, He H, Li J, Chen W, Wang X, Guo L, et al. Genome-Wide Analysis of DNA Methylation and Gene Expression Changes in Two Arabidopsis Ecotypes and Their Reciprocal Hybrids. Plant Cell. 2012;24: 875–892. Available: http://www.plantcell.org/cgi/doi/10.1105/tpc.111.094870

15. Greaves IK, Groszmann M, Ying H, Taylor JM, Peacock WJ, Dennis ES. Trans chromosomal methylation in Arabidopsis hybrids. Proc Natl Acad Sci U S A. 2012;109: 3570–3575. doi:10.1073/pnas.1201043109

16. Greaves IK, Groszmann M, Wang A, Peacock WJ, Dennis ES. Inheritance of Trans Chromosomal Methylation patterns from Arabidopsis F1 hybrids. Proc Natl Acad Sci U S A. 2014;111: 2017–2022. doi:10.1073/pnas.1323656111

17. Johannes F, Porcher E, Teixeira FK, Saliba-Colombani V, Simon M, Agier N, et al. Assessing the impact of transgenerational epigenetic variation on complex traits. PLoS Genet. 2009;5. doi:10.1371/journal.pgen.1000530

18. Vongs A, Kakutani T, Martienssen RA, Richards EJ. Arabidopsis thaliana DNA methylation mutants. Science. 1993;260: 1926–1928. doi:10.1126/science.8316832

19. Zemach A, Kim MY, Hsieh P-H, Coleman-Derr D, Eshed-Williams L, Thao K, et al. The Arabidopsis Nucleosome Remodeler DDM1 Allows DNA Methyltransferases to Access H1-Containing Heterochromatin. Cell; 2013;153: 193–205. Available: http://dx.doi.org/10.1016/j.cell.2013.02.033

20. Colome-Tatche M, Cortijo S, Wardenaar R, Morgado L, Lahouze B, Sarazin A, et al. Features of the Arabidopsis recombination landscape resulting from the combined loss of sequence variation and DNA methylation. Proc Natl Acad Sci. National Acad Sciences; 2012;109: 16240–16245. Available: http://www.pnas.org/content/109/40/16240.short

21. Cortijo S, Wardenaar R, Colome-Tatche M, Gilly A, Etcheverry M, Labadie K, et al. Mapping the epigenetic basis of complex traits. Science. 2014;343: 1145–1148. doi:10.1126/science.1248127

22. Soppe WJ, Jacobsen SE, Alonso-Blanco C, Jackson JP, Kakutani T, Koornneef M, et al. The late flowering phenotype of fwa mutants is caused by gain-of-function epigenetic alleles of a homeodomain gene. Mol Cell. 2000;6: 791–802. Available: http://www.ncbi.nlm.nih.gov/pubmed/11090618

23. Groszmann M, Gonzalez-Bayon R, Greaves IK, Wang L, Huen AK, Peacock WJ, et al. Intraspecific Arabidopsis hybrids show different patterns of heterosis despite the close relatedness of the parental genomes. Plant Physiol. 2014;166: 265–280. doi:10.1104/pp.114.243998

24. Wang L, Greaves IK, Groszmann M, Wu LM, Dennis ES, Peacock WJ. Hybrid mimics and hybrid vigor in Arabidopsis. Proc Natl Acad Sci. 2015;112: E4959–E4967. doi:10.1073/pnas.1514190112

25. Bennett E, Roberts J a., Wagstaff C. Manipulating resource allocation in plants. J Exp Bot. 2012;63: 3391–3400. doi:10.1093/jxb/err442

26. Massonnet C, Vile D, Fabre J, Hannah M a., Caldana C, Lisec J, et al. Probing the Reproducibility of Leaf Growth and Molecular Phenotypes: A Comparison of Three Arabidopsis Accessions Cultivated in Ten Laboratories. Plant Physiol. 2010;152: 2142–2157. doi:10.1104/pp.109.148338

27. Tsukahara S, Kobayashi A, Kawabe A, Mathieu O, Miura A, Kakutani T. Bursts of retrotransposition reproduced in Arabidopsis. Nature; 2009;461: 423–427. doi:10.1038/nature08351

28. Kooke R, Johannes F, Wardenaar R, Becker F, Etcheverry M, Colot V, et al. Epigenetic Basis of Morphological Variation and Phenotypic Plasticity in Arabidopsis thaliana. Plant Cell. 2015;27: 337–348. doi:10.1105/tpc.114.133025

29. Rosado A, Li R, van de Ven W, Hsu E, Raikhel N V. Arabidopsis ribosomal proteins control developmental programs through translational regulation of auxin response factors. Proc Natl Acad Sci U S A. 2012;109: 19537–19544. doi:10.1073/pnas.1214774109

30. Pinon V, Etchells JP, Rossignol P, Collier S a, Arroyo JM, Martienssen R a, et al. Three PIGGYBACK genes that specifically influence leaf patterning encode ribosomal proteins. Development. 2008;135: 1315–1324. doi:10.1242/dev.016469

31. Gachomo EW, Jimenez-Lopez JC, Baptiste LJ, Kotchoni SO. GIGANTUS1 (GTS1), a member of Transducin/WD40 protein superfamily, controls seed germination, growth and biomass accumulation through ribosome-biogenesis protein interactions in Arabidopsis thaliana. BMC Plant Biol.; 2014;14: 1–17. doi:10.1186/1471-2229-14-37

32. Oikawa K, Kasahara M, Kiyosue T, Kagawa T, Suetsugu N. CHLOROPLAST UNUSUAL POSITIONING1 Is Essential for proper Chloroplast positioning. The Plant Cell, 2003;15: 2805–2815. doi:10.1105/tpc.016428.tions.

33. Dapp M, Reinders J, Bediee A, Balsera C, Bucher E, Theiler G, et al. Heterosis and inbreeding depression of epigenetic Arabidopsis hybrids. Nat Plants. 2015;1: 1–8. doi:10.1038/nplants.2015.92

34. Stroud H, Do T, Du J, Zhong X, Feng S, Johnson L, et al. Non-CG methylation patterns shape the epigenetic landscape in Arabidopsis. Nat Struct Mol Biol. 2014;21: 64–72. doi:10.1038/nsmb.2735

35. Meyer RC. Heterosis of Biomass Production in Arabidopsis. Establishment during Early Development. PLANT Physiol. 2004;134: 1813–1823. Available: http://www.plantphysiol.org/cgi/doi/10.1104/pp.103.033001

